# Evaluation of enzyme activity predictions for variants of unknown significance in Arylsulfatase A

**DOI:** 10.1101/2024.05.16.594558

**Authors:** Shantanu Jain, Marena Trinidad, Thanh Binh Nguyen, Kaiya Jones, Santiago Diaz Neto, Fang Ge, Ailin Glagovsky, Cameron Jones, Giankaleb Moran, Boqi Wang, Kobra Rahimi, Sümeyra Zeynep Çalıcı, Luis R. Cedillo, Silvia Berardelli, Buse Özden, Ken Chen, Panagiotis Katsonis, Amanda Williams, Olivier Lichtarge, Sadhna Rana, Swatantra Pradhan, Rajgopal Srinivasan, Rakshanda Sajeed, Dinesh Joshi, Eshel Faraggi, Robert Jernigan, Andrzej Kloczkowski, Jierui Xu, Zigang Song, Selen Özkan, Natàlia Padilla, Xavier de la Cruz, Rocio Acuna-Hidalgo, Andrea Grafmüller, Laura T. Jiménez Barrón, Matteo Manfredi, Castrense Savojardo, Giulia Babbi, Pier Luigi Martelli, Rita Casadio, Yuanfei Sun, Shaowen Zhu, Yang Shen, Fabrizio Pucci, Marianne Rooman, Gabriel Cia, Daniele Raimondi, Pauline Hermans, Sofia Kwee, Ella Chen, Courtney Astore, Akash Kamandula, Vikas Pejaver, Rashika Ramola, Michelle Velyunskiy, Daniel Zeiberg, Reet Mishra, Teague Sterling, Jennifer L. Goldstein, Jose Lugo-Martinez, Sufyan Kazi, Sindy Li, Kinsey Long, Steven E. Brenner, Constantina Bakolitsa, Predrag Radivojac, Dean Suhr, Teryn Suhr, Wyatt T. Clark

## Abstract

Continued advances in variant effect prediction are necessary to demonstrate the ability of machine learning methods to accurately determine the clinical impact of variants of unknown significance (VUS). Towards this goal, the ARSA Critical Assessment of Genome Interpretation (CAGI) challenge was designed to characterize progress by utilizing 219 experimentally assayed missense VUS in the *Arylsulfatase A* (*ARSA*) gene to assess the performance of community-submitted predictions of variant functional effects. The challenge involved 15 teams, and evaluated additional predictions from established and recently released models. Notably, a model developed by participants of a genetics and coding bootcamp, trained with standard machine-learning tools in Python, demonstrated superior performance among submissions. Furthermore, the study observed that state-of-the-art deep learning methods provided small but statistically significant improvement in predictive performance compared to less elaborate techniques. These findings underscore the utility of variant effect prediction, and the potential for models trained with modest resources to accurately classify VUS in genetic and clinical research.

## 1 Introduction

The characterization of variants of unknown significance (VUS) is critical for genetic diagnosis (Richards et al, 2015), newborn screening (Hong et al, 2021; Stark and Scott, 2023), estimating disease burden (Clark et al, 2018; Borges et al, 2020; Chen et al, 2023), and gaining insights into the molecular mechanisms of human genetic disease (Rost et al, 2016; Shendure et al, 2019; Estrada et al, 2021). However, despite their biomedical relevance and need for clinical deployment, sensitive, cost-effective, experimental techniques for characterizing VUS remain elusive. *In silico* predictors hold such promises for characterizing VUS, but have yet to share in the recent performance increases experienced by protein structure prediction (Kryshtafovych et al, 2021).

Towards the understanding and improvement of *in silico* predictors of variant functional effect, the Critical Assessment for Genome Interpretation (CAGI) organization has continually facilitated the use of experimentally derived and real-world data to train and evaluate state-of-the-art predictive tools (The Critical Assessment of Genome Interpretation Consortium, 2024). As part of the sixth round of CAGI challenges (CAGI6), a blinded competition was conducted to assess the community’s performance at predicting the impact of missense variants in the Arylsulfatase A gene (*ARSA*) on its enzymatic activity.

Bi-allelic variants in *ARSA* can lead to Metachromatic Leukodystrophy (MLD), an autosomal-recessive lysosomal storage disorder characterized by neuro-cognitive decline, and variable age of onset and severity (Van Rappard et al, 2015). Without early intervention, patients with the most severe, Late-Infantile, form of the disease survive only into early childhood, whereas Adult onset patients may not be diagnosed until the fifth decade of life, and, in some cases, have been mistaken for having Alzheimer’s disease (Martin et al, 2013; Johannsen et al, 2001; Stoeck et al, 2016). Multiple studies have highlighted a strong genotype-phenotype relationship in MLD when considering well-characterized and common pathogenic *ARSA* mutations (Kappler et al, 1991; Trinidad et al, 2023). For these reasons, understanding the performance of *in silico* methods at assessing *ARSA* variant impact is particularly important. As more novel variants are identified, understanding their potential contribution to disease will be a high priority to the newborn screening community so that early and effective therapeutic responses to disease can be employed (Hong et al, 2021).

An evaluation dataset of *ARSA* variants was previously tested for their impact on enzymatic activity using a tandem mass spectrometry assay (Trinidad et al, 2023). Variants were curated from patients reported in the literature, collected by the MLD Foundation, or reported in databases of population variants such as gnomAD (Karczewski et al, 2020). For the *ARSA* Challenge, predictions of *ARSA* variant impact, represented as the resulting protein’s enzymatic activity relative to the wild-type (%WT activity) were solicited before the experimentally derived %WT activity values were published.

In this paper, we report the findings of the *ARSA* Challenge. We observed that top predictor performance was consistent with previous, similarly designed challenges (Clark et al, 2019). Also, consistent with previous challenges, we observed that models employing disparate methodologies and training data were still highly correlated with each other, suggesting common underlying characteristics influencing predictor performance. Finally, we observed that simple machine-learning models performed on par with advanced deep-learning methods, suggesting that feature engineering and quality of the training data can compensate for the complexity of the learning algorithm used in the model (Nguyen, 2024).

## 2 Results

### 2.1 Challenge design and participation

The ARSA CAGI challenge was structured similar to previous related challenges (Clark et al, 2018, 2019). First, a *Curated* set of 274 ARSA single-nucleotide-variants (SNVs), observed in the population or of known disease significance was provided prior to publication of experimentally assayed %WT activity values (Trinidad et al, 2023).

Variants were released to the community through the CAGI organization website, requesting participants to provide predicted %WT enzymatic activity values associated with each missense mutation in the ARSA protein (ENSP00000216124). Prediction submissions were accepted from July 15, 2022 to November 15, 2022. Additional publicly available predictions from published models were also collected, but were evaluated separately from submitted predictions when an overall ranking of models was generated (Adzhubei et al, 2010; Ioannidis et al, 2016; Pejaver et al, 2020; Cheng et al, 2023).

Fifteen teams participated in the challenge, submitting predictions form a total of 65 predictors (Table 2, Table 1). Four teams, comprised mostly undergraduate participants (but also high school and graduate students), of a two-week genetics and coding bootcamp that was synchronized with the challenge timeline and academic calendars. Bootcamp participants hailed from 8 different countries: Argentina, Australia, China, France, Italy, Japan, Turkey, and the United States, including Puerto Rico.

**Table 1:** A table of each predictor, its primary reference if available, categories of features used, and training data sources.

| Method name | Reference | PolyPhen, SIFT, Provean or other features from dbNSFP | Structure Based Features | PSSM, MSA Based Features | ML Method | Training Database |
| --- | --- | --- | --- | --- | --- | --- |
| Bootcamp Team 1 | NA | Yes | No | No | AdaBoost | ClinVar |
| Bootcamp Team 2 | NA | Yes | No | No |  | ClinVar |
| Bootcamp Team 3 | Nguyen (2024) | Yes | No | No | Random forest | ClinVar |
| Bootcamp Team 4 | NA | Yes | No | No |  | ClinVar |
| Ken Chen | NA | No | No | No | Neural Network | ClinVar |
| Meta-EA clinical | Katsonis and Lichtarge (2014) | Yes | No | Yes | None | None |
| TCS Research India | NA | No | No | No | Neural Network | humsavar, Clinvar, gnomAD v3.1.2, the Pompe disease mutation database, the Leiden Open Variation Database 3.0 |
| Shoni.cagi6 | NA | No | No | No | Hybrid, GENN | ClinVar |
| Random Forest Ensemble | NA | Yes | Yes | Yes | Random Forest Ensemble | ClinVar |
| Qafi | Özkan et al (2021) | Yes | Yes | Yes | Multiple Linear Regression and Ensemble | Deep mutational scanning data for 30 proteins |
| Evolutionary Index | Frazer et al (2021) | No | No | Yes | Neural Network | MAVEDB |
| E-SNPs & GO | Manfredi et al (2022) | No | No | No | SVM | ClinVar, humsavar |
| Zero-shot protein language | Sun and Shen (2023) | No | No | No | Transformer encoder/decoder | Pfam |
| SNPMuSiC | Ancien et al (2018) | Yes | Yes | No | Artificial Neural Network | ClinVar |
| Random Forest Labeling | NA | Yes | No | No | Random Forest | ClinVar |

**Table 2:**
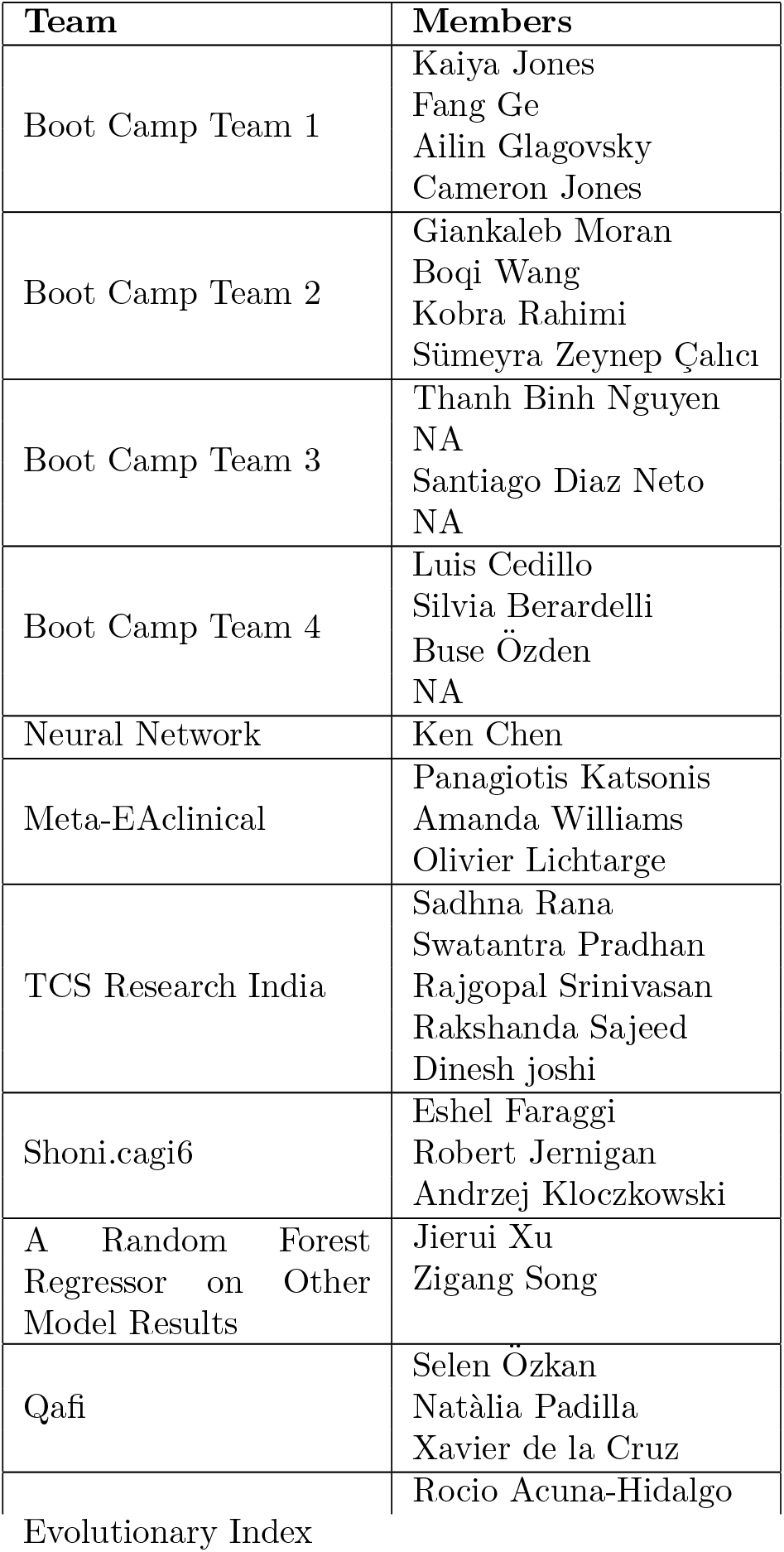
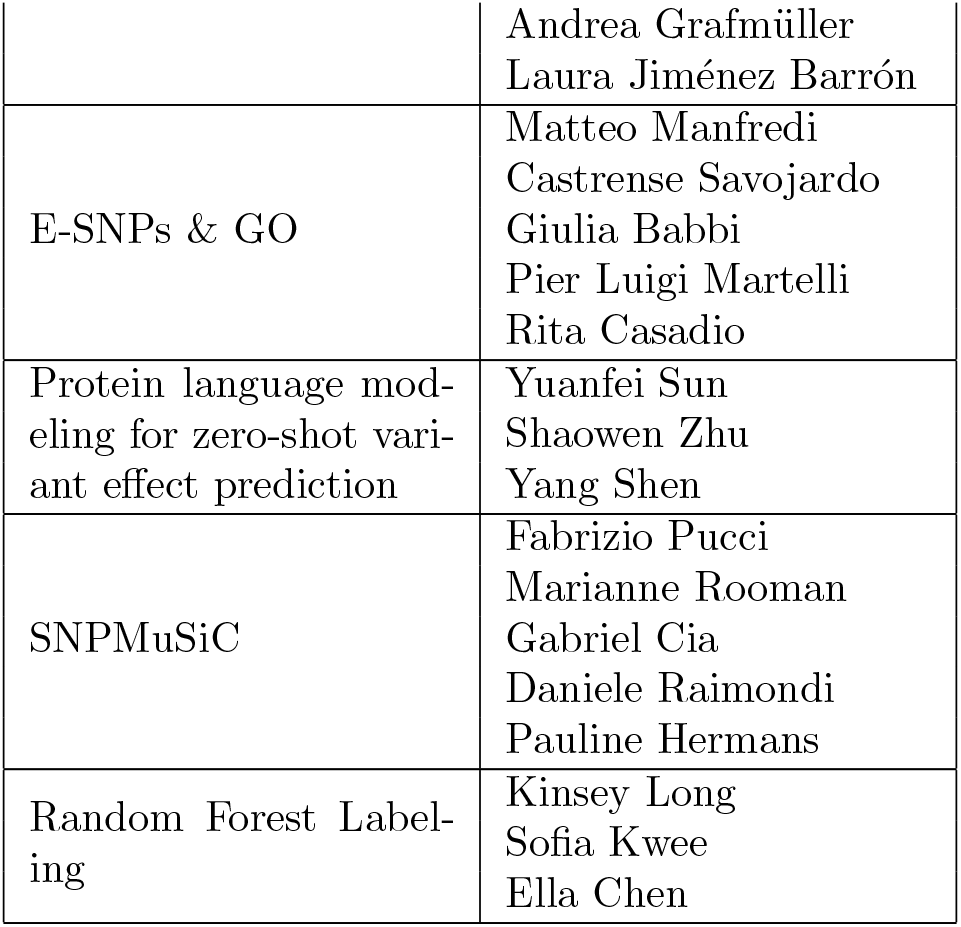
A table of each predictor and individuals who participated in the model’s development or submission to the ARSA CAGI Challenge.

### 2.2 Performance at predicting enzymatic activity values

To evaluate model performance, we calculated the relative ranking of each predictor according to Pearson’s correlation, Kendall’s tau, area under the receiver operating characteristic curve (AUC), and truncated AUC (Section 3.2). Pearson’s correlation and Kendall’s tau are used for predicting the experimentally measured percent wild-type activity (%WT) as a continuous value. AUC and truncated AUC are used for classifying variants as pathogenic or benign, where the variants were considered pathogenic if the %WT was ≤13%. We look at truncated AUC in addition to AUC because a region of low false positive rate (FPR) is of particular interest in clinical settings. A final, overall ranking was achieved by then taking the average rank according to these four metrics (Fig. 1, Fig. 2). Since a team could have multiple predictors, we first performed ranking within each team and picked the best ranking predictor as its representative in the final overall ranking. Assessors were blinded to model names until models were ranked and the best ranking predictor for each team was chosen. A mapping between original identifiers and representative model names can be found in Table S5.

**Fig. 1:**
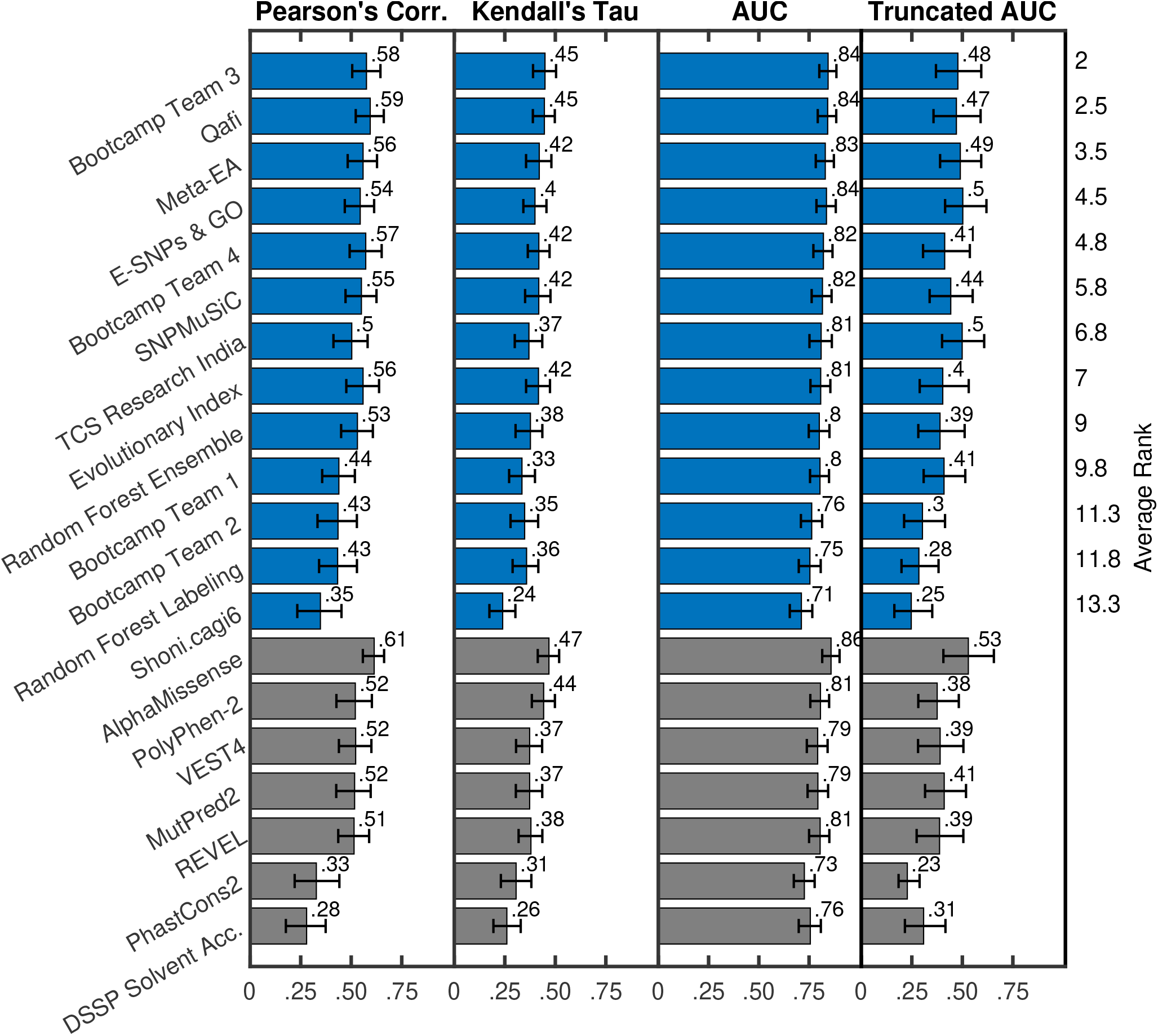
Model performance based on key metrics: Model performance based on Pearson’s correlation, Kendall’s tau, AUC, and Truncated AUC; along with bootstrap-based 90% confidence intervals. The best-performing model for each team is shown in blue. Baseline models and individual feature (evolutionary and structure) based performance are shown in grey. Models are sorted on the y-axis based on their average ranking according to the four metrics used. Only models with Pearson’s correlation greater than 0.25 are shown in the figure (13 out of 15 submitted models, five out of five baseline models, and two out of seven fundamental features). For teams that submitted multiple models, we show the performance of their best model. Figure S1 in Supplementary Materials shows performance for all models (or the best models for teams that submitted more than one).

**Fig. 2:**
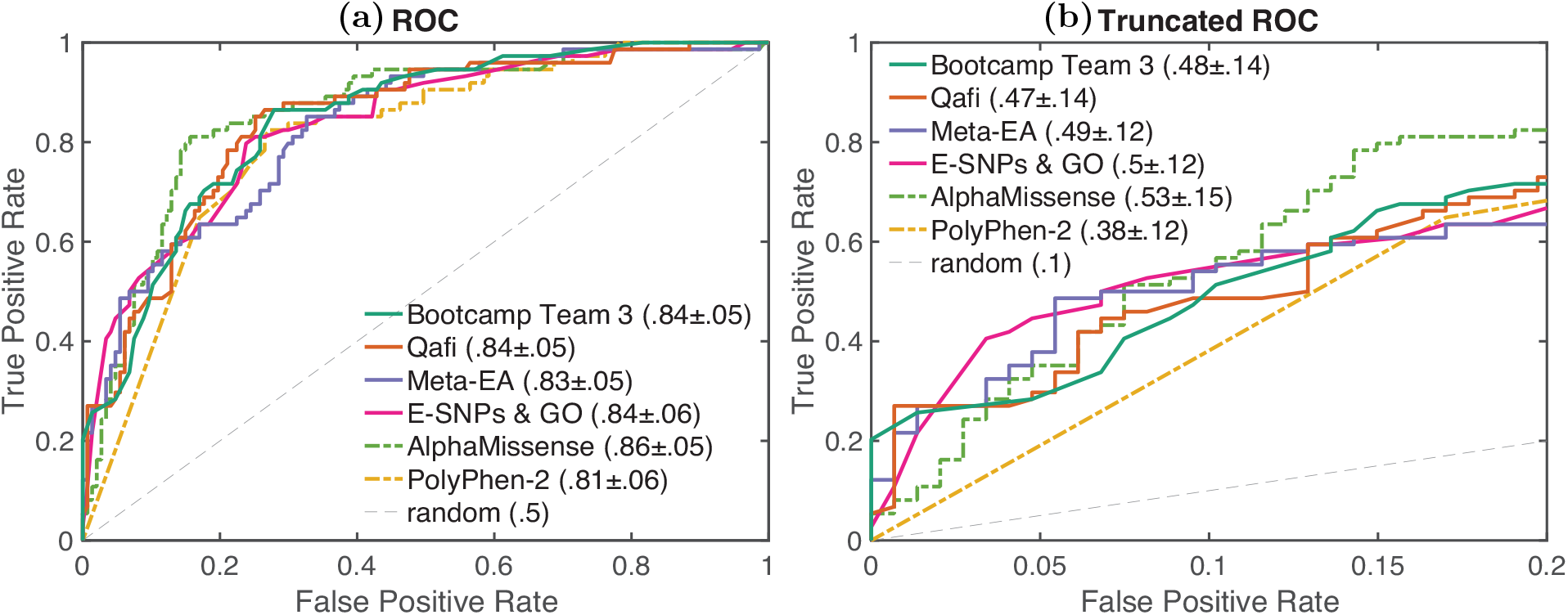
**a)** ROC and **b)** Truncated ROC: The receiver operating characteristic (ROC) and truncated ROC curves for the best-performing model for each team and baselines.AUC and Truncated AUC values are shown along with 1.96*×*standard deviation from their bootstrap estimates. Results are shown for the top four submitted models and top two baseline models (based on their average ranking according to the four metrics used).

Out of all ARSA challenge participants, we observed that Bootcamp Team 3’s model performed the best (described in Section 3.5), achieving an average rank of 2 across the four evaluation metrics considered (Pearson’s Corr = 0.576, rank = 2; Kendall’s Tau = 0.449, rank = 1; AUC = 0.845, rank = 1; Truncated AUC = 0.478, rank = 4) (Nguyen, 2024). Qafi was the second best performing model with an average rank of 2.5 across the four evaluation metrics (Pearson’s Corr = 0.594, rank = 1; Kendall’s Tau = 0.446, rank = 2; AUC = 0.843, rank = 2; Truncated AUC = 0.470, rank = 5) (Özkan et al, 2021).

#### 2.2.1 Comparison with publicly available tools

We also evaluated the performance of five publicly available predictors: PolyPhen-2 (Adzhubei et al, 2010), VEST4 (Carter et al, 2013), REVEL (Ioannidis et al, 2016), MutPred2 (Pejaver et al, 2020), and AlphaMis-sense (Cheng et al, 2023); grey bars in Fig. 1, Fig. 2). Ranking was performed separately for this group of models.

Out of these models, AlphaMissense showed the highest performance in evaluation (Pearson’s Corr = 0.614, rank = 1; Kendall’s Tau = 0.468, rank = 1; AUC = 0.859, rank = 1; Truncated AUC = 0.530, rank = 1. Furthermore, AlphaMissense was the only publicly available model to perform better than the top-performing challenge participant (Bootcamp Team 3). Though the difference in performance was statistically significant on all four metrics, the effect size was relatively small. Statistical significance was determined using a one-sided binomial test with number of wins on 1000 bootstrap samples as the test statistic. AlphaMissense won 875, 763, 760 and 755 times on Pearson’s Corr., Kendall’s Tau, AUC and Truncated AUC, giving p-values less than 10^−139^, 10^−65^, 10^−63^ and 10^−61^, respectively. The p-value was computed as the probability that the Binomial(0.5, 1000) variable is greater than or equal to the number of wins.

### 2.2.2 Comparison with fundamental features

In addition to comparing participant performance with publicly available state-of-the-art predictors, we also selected individual features based on evolutionary conservation (PhastCons2), biochemical properties (Grantham scores), and protein structure characteristics (solvent accessibility, and backbone kappa, alpha, phi, and psi values using the PDB structure 1AUK) for evaluation (Siepel et al, 2005; Grantham, 1974; Kabsch and Sander, 1983).

PhastCons2 (Pearson’s Corr = 0.327; Kendall’s Tau = 0.306; AUC = 0.727; Truncated AUC = 0.228) and solvent accessibility (Pearson’s Corr = 0.279; Kendall’s Tau = 0.260; AUC = 0.756; Truncated AUC = 0.308) were the best two performing features. Other features (Grantham scores and backbone’s dihedral angles) showed poor correlation (Pearson’s Corr ≤ 0.25) with the %WT activity values and were excluded from the figure. Figure S1 in Supplementary material shows performance for all fundamental features.

### 2.2 Correlation between predictors

We measured the correlation of predictions between models (Fig. 3). Consistent with previous challenges, models were all more correlated with each other than with %WT activity values (Clark et al, 2019). The top 3 performing methods in term of Pearson’s correlation (AlphaMissense, Quafi, and Bootcamp Team 3) achieved correlation with %WT activity of less than 0.60, but were more correlated with each other (Pearson’s correlations *>* 0.76. This high correlation could be the result of shared methodologies or training features. For example, Qafi, Meta-EA and BC Team 3 all utilize REVEL as a feature, while the other top-scoring predictors do not (Table 1). Qafi and Meta-EA also both employ PSSM/MSA-based features. Generally, Qafi had higher correlations with other models across the board (0.74 − 0.93), likely due to being an ensemble method that incorporated features of variant impact predictors, structural information, and PSSM/MSA data. Bootcamp team models all utilized the same training features, so high correlation between them is anticipated. SNP-MuSiC, a predominantly structure-based method, showed lower correlation across the board compared to other submitters, indicating a more unique methodology. With regards to training data, all of the top performing models except for Qafi and Meta-EA (a non-machine learning model) were trained on either ClinVar or Humsavar databases, likely also leading to stronger correlation between them (Table 1).

**Fig. 3:**
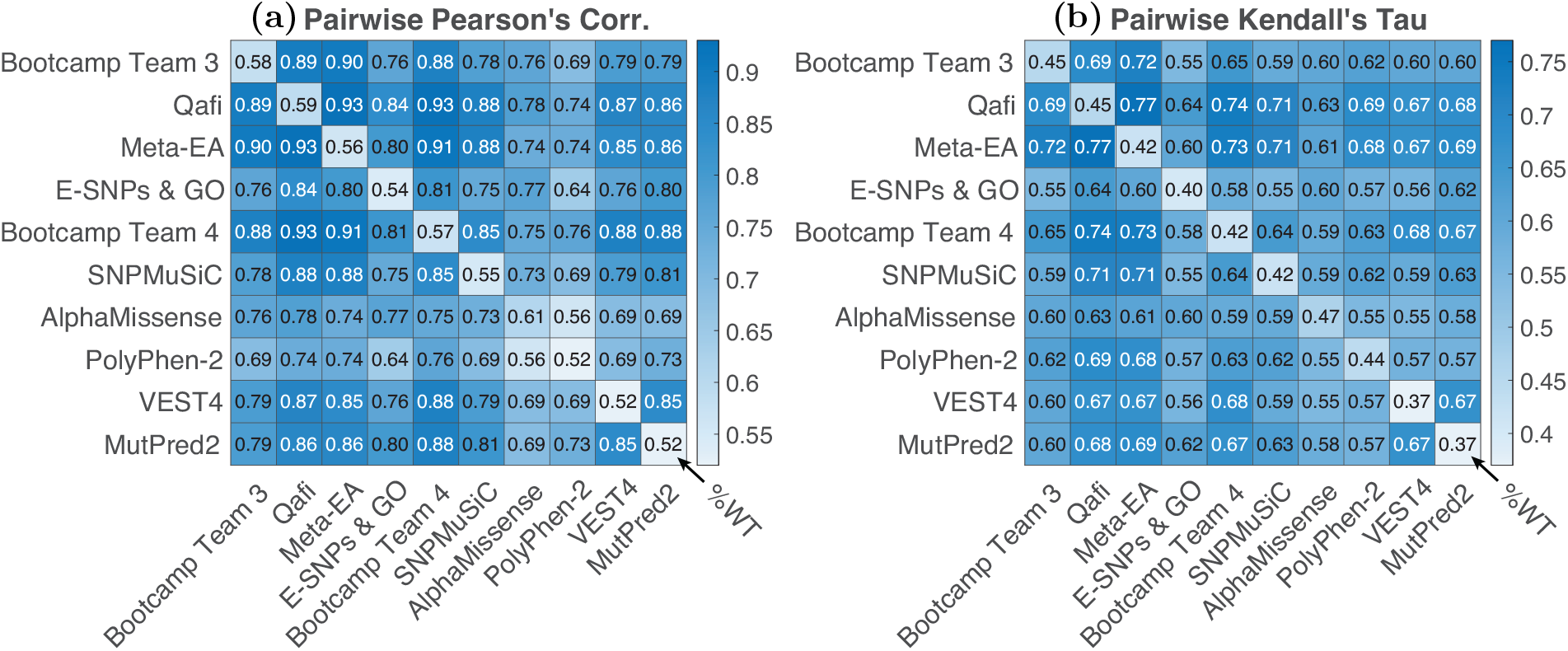
Pairwise correlation: pairwise correlation coefficients between models, with Pearson’s correlation coefficients shown on the left **(a)** and Kendall’s Tau shown on the right **(b)**. The diagonal for each figure shows the correlation of that model with observed enzymatic activity values. Results are shown for the top six submitted models and top four baseline models (based on their average ranking according to the four metrics used).

### 2.4 Difficult-to-predict variants

We considered the hardest-to-predict pathogenic and benign variants to search for trends (Section 3.6). For the current analysis we consider 13% WT activity to be the threshold that distinguishes between pathogenic and benign variants, and considered all mutations, not just those in the evaluation dataset. We found that the top 10 difficult-for-all pathogenic variants had a median %WT activity of 1.28, making them closer to the pathogenic-benign threshold (13 %WT activity) on average compared to the other pathogenic variants, suggesting that predictors have more trouble with hypomorphic mutations with residual activity (Fig. 4 (a)) (Muller, 1932). Out of all pathogenic variants, 67% have a lower %WT activity (a higher pathogenic effect). The top 10 difficult-for-all benign variants with a median 27.6 %WT activity are also closer to the pathogenic-benign threshold on average compared to the other benign variants. Out of all benign variants, 77% have a higher %WT activity (a higher benign effect).

**Fig. 4:**
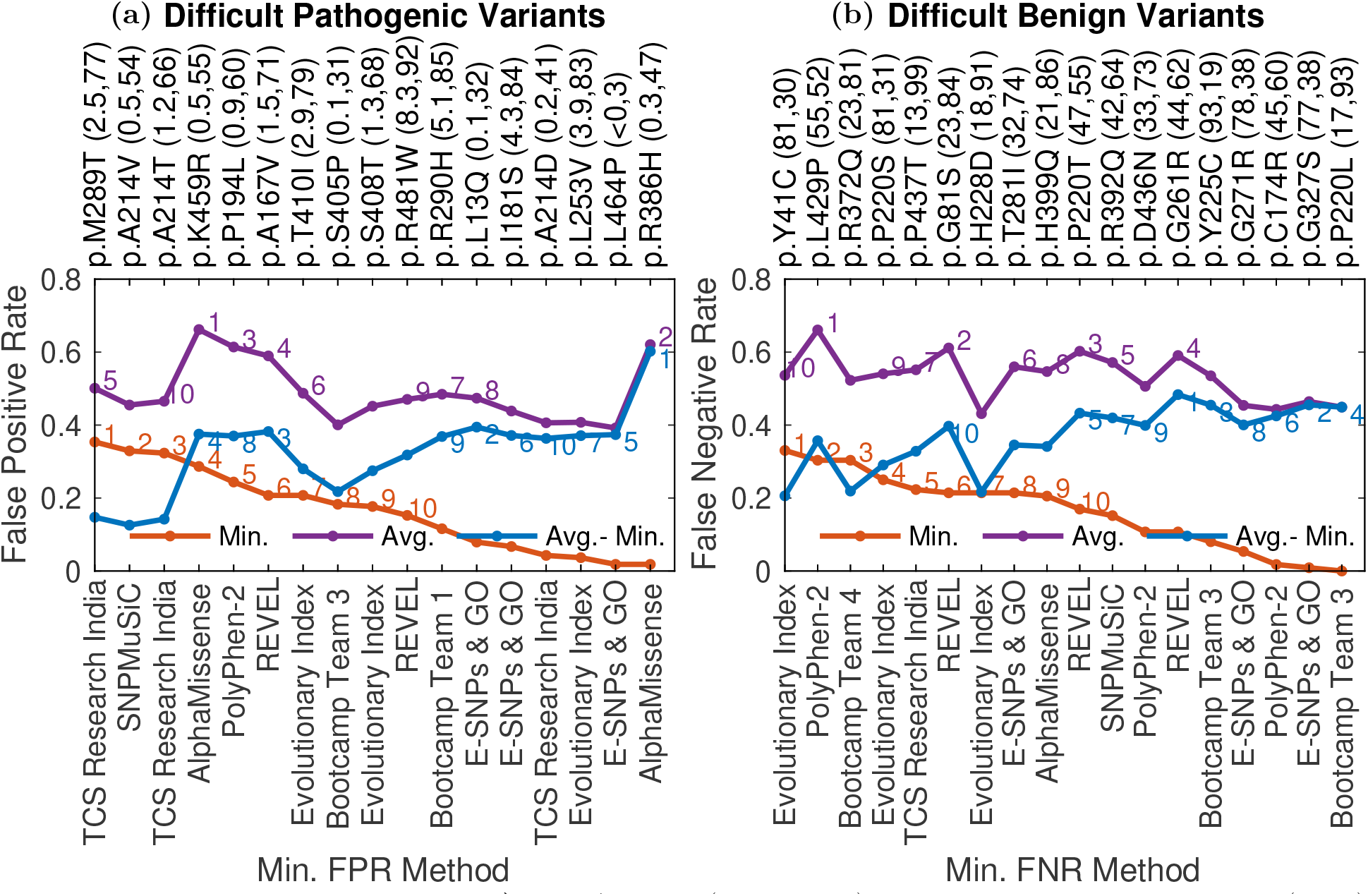
Difficult to classify variants: **a)** The Average (purple line), Minimum False Positive Rate (FPR) (red line), along with their difference (blue line), is given for the difficult to predict pathogenic variants. FPR for each pathogenic variant-predictor pair was calculated as the proportion of all benign variants that are scored higher by the method compared to the variant. The %WT activity (rounded to 1 decimal place) for each variant and the percent of all pathogenic variants attaining a lower %WT activity are shown in the parenthesis next to the variant identifier. The methods attaining the minimum FPR for that variant are also shown along the bottom horizontal axis. **b)** Average (purple line) and Minimum False Negative Rate (FNR) (red line) and their difference (blue line) are given for difficult-to-predict benign variants. FNR for each benign variant-predictor pair was calculated as the proportion of all pathogenic variants that are scored lower by the method compared to the variant. The %WT activity (rounded to the closest integer) for each variant and the percent of all benign variants attaining a higher %WT activity are shown in the parenthesis next to the variant identifier. The methods attaining the minimum FNR are shown along the bottom x-axis. Difficult-to-predict mutations were defined as described in Section 3.6.

Variants p.A214V and p.T410I, among the top 10 difficult-for-all pathogenic variants, are established pathogenic variants in ClinVar annotated as P and LP, respectively. The difficulty in predicting the pathogenicity of these variants is surprising since the variants might be present in the training set of the supervised predictors. The variants have a low %WT activity of 0.45 and 2.9, respectively. However, 54% and 79% of pathogenic variant have a lower %WT activity than p.A214V and p.T410I, respectively. This provides a probable explanation for the difficulty in their prediction, especially for p.T410I. Another expla-nation is that these mutations just happen to be false positives in terms of both their %WT activity values and their annotation in ClinVar. Indeed their frequency in patients is low: neither mutation was identified in curated patients in Trinidad et al (2023), and they both have low allele frequencies in gnomAD.

Variants p.R372Q and p.T281I, among the top 10 difficult-for-all benign variants, have known clinical significance in ClinVar, however, they are annotated as P/LP and LP respectively. Furthermore, although in the benign range, they have low %WT activity of 23.3 and 31.9, with 81% and 74% of the benign variants having a higher %WT activity, respectively. The conflicting annotation in ClinVar and the low %WT activity provide a probable explanation for the difficulty in their prediction.

AlphaMissense and E-SNPs & GO attain the minimum FPR multiple times among the top 10 pathogenic variants with highest difference between average and minimum FPR (Gap list variants). Other predictors might gain insights from the two methods to improve their predictions on these variants. REVEL, PolyPhen-2, E-SNPs & GO and Bootcamp Team 3 attain the minimum FNR multiple times among the top 10 benign variants with the highest difference between average and minimum FNR. Other predictors might gain insights from the four methods to improve their predictions on these variants.

## 3 Methods

### 3.1 Evaluation set

An *Evaluation* set of 219 ARSA variants was obtained as a subset of the *Curated* set by removing variants with a definitive classification in the December, 2022, release of ClinVar (Landrum et al, 2018). This version of ClinVar represents the release of the database immediately following the challenge’s prediction submission deadline (Nov. 15, 2022). Only the ClinVar variants with clinical significance *Pathogenic, Likely pathogenic, Pathogenic/Likely pathogenic, Benign, Likely benign* or *Benign/Likely benign* without annotation of *Conflicting interpretations* in the review status were removed from the *Curated* set.

All variants with activity levels below 13%WT activity were considered to be pathogenic (Evaluation set: 73, Curated set: 111), as per previously defined threshold (Supplementary Table S1) (Trinidad et al, 2023). Those mutations with greater than 13% WT were considered benign (Evaluation set: 146, Curated set: 163). No mutation had an overall allele frequency in gnomAD (version 2.0.1) greater than 1%, though p.P220L had an allele frequency of 2.7% in the Finnish ancestry group within gnomAD.

### 3.2 Evaluation metrics

To evaluate the predictors, we used standard and clinically relevant performance metrics. Two general classes of metrics were considered: (1) those that treat measured %WT activity as a continuous value, and (2) binary classification metrics that treat each mutation as either pathogenic or benign.

Let *n* be the total number of variants in the evaluation set. Let *y*_*i*_ and *ŷ*_*i*_ denote the experimentally measured %WT activity and its prediction/score, respectively, for variant *i* in the evaluation set. We transform the original predictions from a method, baseline or fundamental feature for a meaningful evaluation, as discussed in Section 3.3. Let *z*_*i*_ = I[*y*_*i*_ *<* 13] be the ground truth pathogenicity class label for variant *i*, taking value 1 and 0 for a pathogenic and benign variant, respectively. Let 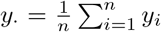 and 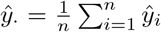 be the average experimental and predicted %WT activity, respectively, over the Evaluation set.

#### 3.2.1 %WT activity evaluation metrics

For the regression task of predicting the percent wild-type activity (%WT), we incorporated Pearson’s correlation (*r*) and Kendall’s Tau (*τ*) as our main two metrics and R-Squared (*R*^2^) as an additional metric; see Supplementary Table S2. The three metrics are defined, as per their standard definitions, below.

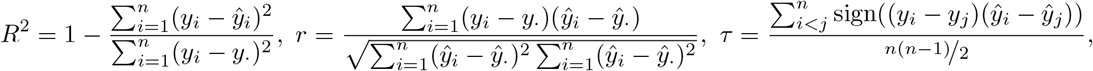

where sign(*x*) is −1, 0 or 1, when *x <* 0, *x* = 0 or *x >* 0, respectively. *R*^2^, a value in the range [−∞, 1], is used to measure the fraction of variance of %WT activity that is explained by the predictor. A predictor with 0 mean square error has a perfect *R*^2^ of 1. An *R*^2^ of 0 indicates that the predictor is only as good as the trivial predictor given by *y*_*·*_ and a negative *R*^2^ would indicate an even worse performance. *R*^2^ is the most stringent metric considered in our evaluation which requires a predictor to be well calibrated. Thus we do not use it for ranking the methods. Pearson’s correlation, a value in the range [−1, 1], is used to measure the statistical association between %WT activity and the predictor assuming a linear relationship between them. Values closer to 1 would indicate a strong correlation, whereas that near or equal to 0 would indicate a weak or no correlation. A negative value would indicate an inverse association between %WT activity and the predictions. Kendall’s Tau, a value in the range [−1, 1], is used to measure the association between %WT activity and its predictions, non-parametrically without assuming a linear relationship, as the difference between the proportions of concordant (pairs of variants ranked correctly by the predictor) and discordant pairs of variants in the Evaluation set. Like Pearson’s correlation, values of 1, −1 and 0 indicate strong positive, negative and no association, respectively.

### 3.2.2 Binary classification metrics

For the classification task of separating pathogenic and benign variants we incorporated the standard binary classification metrics of the Receiver Operating Characteristic (ROC) curve, area under the ROC curve (AUC) and recently derived clinically relevant metrics of area under the truncated ROC curve (tAUC), local likelihood ratio plus (lr^+^) and posterior probability of pathogenicity (*ρ*) curves. Additionally, we provide the proportion of variants for which a predictor provides supporting, moderate, and strong levels of evidence and the corresponding False Discovery Rates (FDR). We define all the classification measures next.

The ROC curve is generated by plotting the True Positive Rate (TPR) against the False Positive Rate (FPR) for the entire range of classification thresholds. TPR and FPR are defined as the proportion of correctly classified pathogenic and incorrectly classified benign variants, respectively; i.e., for a given threshold *t*, below (above) which variants are predicted to be positive (negative), TPR (FPR) is defined as the proportion of pathogenic (benign) variants that have predicted %WT activity smaller than or equal to *t*. Precisely,

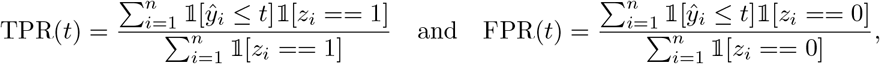

where 1 is the indicator function equal to 1 when the condition within the bracket is satisfied, otherwise it is 0. Note that we define TPR and FPR with ≤ *t*, instead of ≥ *t*, because *ŷ*_*i*_ is a prediction of %WT activity, where a high value corresponds to pathogenicity, unlike the typical classification score. To construct the ROC curve for a predictor, TPR(*t*) is plotted against FPR(*t*) as *t* is varied taking values among its %WT activity predictions, starting from the smallest to the largest. Ties among the predictor scores require special consideration for correctly computing and plotting the ROC curve (Fawcett, 2006). AUC is numerically computed as the area under the ROC curve, requiring special considerations for ties (Fawcett, 2006). Mathematically, AUC is the probability that a random pathogenic variant has a predicted %WT activity smaller than that of a random benign variant. For a more accurate definition that accounts for ties in predictions, a term giving the probability of randomly picked pathogenic and benign variants having equal predictions is added after multiplying by ½ (Byrne, 2016). Intuitively, AUC captures how well a predictor separates the pathogenic and benign variant distributions. AUC takes value in the range from 0 to 1. A value above 0.5 is expected from any classifier that captures some signal for pathogenicity. A salient aspect of AUC is that it is insensitive to class imbalance or the proportion of pathogenic and benign variants in the dataset (Hanley and McNeil, 1982).

Though AUC is a useful measure for the overall classification performance, it has limitations when applied to a decision-making setting such as the one encountered in the clinic. Typically, clinically relevant score thresholds that determine the variants satisfying Supporting, Moderate or Strong (Richards et al, 2015) evidence lie in a region of low false positive rate (FPR). A measure well-suited to capture the clinical significance of a predictor ought to be sensitive to the variations in the classifier’s performance in the low FPR region (when predicting pathogenicity). However, the contribution of the low FPR region to AUC is relatively small. To mitigate this problem, we incorporate the recently derived Truncated AUC (tAUC), the area under the ROC curve truncated to the [0, 0.2] FPR interval (The Critical Assessment of Genome Interpretation Consortium, 2024). The truncated AUC is normalized to span the entire [0, 1] range by dividing it by 0.2, the maximum possible area under the un-normalized truncated ROC.

### 3.3 Prediction normalization and missing values

We noticed that some teams submitted predictions for posterior probability of pathogenicity instead of %WT activity. The baseline methods considered in this work also predict the posterior probability of pathogenicity. Directly applying the evaluation metrics on such predictions would give an incorrect assessment of their performance, since posterior probability of pathogenicity and %WT activity are anti-correlated. Furthermore, even the teams that indeed submit predictions for %WT activity, use a different scale than the experimentally measured %WT activity values. The measured values range from the minimum of −0.016 to the maximum of 161.17, whereas teams were expected to submit predictions on a scale, where 0 means no activity, 1 means %WT activity and values *>* 1 mean higher than %WT activity, as per the submission format. For a meaningful interpretation of computed R-Squared, it is necessary that the predictions are on the same scale as the %WT activity values.

To address this issue, we transform the predictions for %WT activity as *ŷ*_*i*_ = 100*r*_*i*_, where *r*_*i*_ represents the original prediction for variant *i*. We transform the predictions for posterior probability of pathogenicity as *ŷ*_*i*_ = 100(1 − *r*_*i*_). Since we do not always know which methods predict %WT activity and which ones predict posterior probability of pathogenicity, we use AUC computed on the original predictions as a heuristic to separate those cases. Precisely, if using the original predictions give an AUC *<* 0.5, we interpret them to represent posterior probabilities of pathogenicity and use *ŷ*_*i*_ = 100(1 − *r*_*i*_), otherwise we use *ŷ*_*i*_ = 100*r*_*i*_. Note that using *ŷ*_*i*_ = 100*r*_*i*_ would give exactly the same values for Pearson’s correlation, Kendall’s Tau and AUC as the original predictions. In case of *ŷ*_*i*_ = 100(1 − *r*_*i*_), the sign of Pearson’s correlation and Kendall’s Tau is flipped but their magnitude is the same and the transformed AUC is 1− AUC of the original predictions.

The fundamental features are not explicitly derived to predict the %WT activity or the posterior probability of pathogenicity. For a meaningful evaluation, we apply the following transform to the fundamental features. If AUC computed with an un-transformed feature is greater than equal to 0.5 we Use 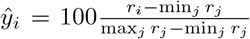 (Min-max normalization followed by multiplication by 100), otherwise we use 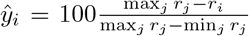 (Min-max normalization on the negated predictions followed by multiplication by 100).

In the case where a submitted method, baseline or a feature had missing values, we performed imputation by the mean of the non-missing predictions. Prediction normalization was performed after the imputation.

### 3.4 Uncertainty quantification

To quantify the uncertainty in performance estimation, we obtained 1000 bootstrap estimates of each metric. Each bootstrap sample was created by sampling *n* = 219 (size of the Evaluation set) variants from the Evaluation set with replacement. Each performance metric was estimated on each of the 1000 bootstrap samples. Confidence interval for each metric was obtained from the 5th and 95th percentile of its bootstrap estimates; see Figure 1 and Table S2. A gaussian approximation based 95% confidence interval was provided with the AUC and Truncated AUC values in Figure 2 as 1.96*×*standard deviation derived from the bootstrap estimates.

### 3.5 Bootcamp participant models

A training dataset for Bootcamp models was generated using ClinVar variants annotated as pathogenic or likely pathogenic as positive training data points, and all others as negative data points. Features for each mutation were generated from predictor scores from the dbNSFP website (Liu et al, 2011). Predictor features consisted of predictions from meta-predictors such as REVEL, essentially making the model a meta-meta predictor.

For Bootcamp Team 3, regression models were trained on eight methods, including: logistic regression, perceptron, support vector machine, K-nearest neighbors, decision tree, random forest, neural network, Gaussian Naive Bayes using default parameters in sklearn (Nguyen, 2024). Five-fold cross-validation with shuffle was applied. The best-performing method from cross-validation was found to be the random forest. A search for the optimal parameters was then conducted, including estimator (100, 200, and 300), max features (5,10,15 and auto), max depth (None, 10, and 20), min samples split (2, 5, 8), and min samples leaf (1, 4, 7). The best parameters were found to be max features of 15, max depth of None, min samples split of 2, and min samples leaf of 1.

### 2.6 Categorization of difficult-to-predict mutations

We split difficult-to-predict mutations into pathogenic (Supplementary Table S3) and benign (Supplementary Table S4), where pathogenic and benign mutations are defined as described in Section 3.1. For each pathogenic variant (those with *<* 13% WT activity) and each predictor, we calculated the false positive rate (FPR) for that variant/predictor combination as the proportions of benign variants that had lower %WT activity than the variant at hand. We then considered the minimum and average FPR for each variant across all predictors that attained an overall AUC *>* 0.8. The 10 variants with the highest minimum FPR were then taken as a list of difficult-for-all methods to classify pathogenic mutations (Difficult-for-all list). The top 10 variants with the highest average FPR comprised the set of variants that are difficult to predict on average (Difficult-on-average list). The top 10 variants with the highest difference between average and minimum FPR give a list of pathogenic variants with the highest performance gap (Gap list).

Similarly, for each benign variant (those with *>* 13% WT activity) and each predictor, we calculated the false negative rate (FNR), or proportion of all pathogenic variants that had higher %WT activity than that particular variant. We then calculated the minimum and average FNR for each variant across all model that attained an AUC *>* 0.8. The top 10 variants with the highest minimum FNR were taken as a list of benign variants that were difficult for all methods to classify (Difficult-for-all list), and the top 10 variants with the highest average FNR comprised the set of variants that are difficult to predict on average (Difficult-on-average list). The top 10 variants with the highest difference between average and minimum FNR give a list of benign variants with the highest performance gap (Gap list).

The union of the variants in the three lists was taken to give a final list of difficult variants, separately for the pathogenic and benign categories.

## 4 Discussion

CAGI has served as a platform for using real-world and experimental data to further our understanding of the molecular causes of human disease (The Critical Assessment of Genome Interpretation Consortium, 2024). The CAGI6 *ARSA* challenge assessed *in silico* models at their ability to predict variant impact on ARSA’s enzymatic activity. We observed that top models exhibited similar performance at predicting ARSA variant impact on %WT activity compared to previous studies (Clark et al, 2019). Also, as previously observed, models exhibited high correlations with each other despite differences in methodologies, suggesting shared training features and data as the underlying cause. Ten out of 18 submitted models utilized ClinVar or Humsavar for their training database, while four other models relied on deep mutational scanning data from an overlapping set of proteins. Many submissions were meta-predictors incorporating scores from similar models, such as REVEL, VEST3, PROVEAN, MutationTaster, and PolyPhen-2. The best-performing model, Bootcamp Team 3’s, was a meta-meta predictor.

Consistent with previous findings (Pejaver et al, 2017), we also observed that variant effect predictors designed to predict pathogenicity as a binary class, can be adapted to predicting %WT activity as a continuous variable, though all models seem to have greater difficulty at identifying hypomorphic mutations. Disagreements between *in vitro* assays and the severity of disease affected by a mutation in patients present an additional study limitation. An example of such a variant is p.P428L, which has repeatedly produced %WT activity values lower than expected based on its behavior in patients (Trinidad et al, 2023).

Perhaps the most notable result is the competitiveness between AlphaMissense and Bootcamp Team 3’s model. Though AlphaMissense presents an elegant solution to the problem of predicting missense variant functional effects, it requires a complex deep-learning architecture and significant resources to train. In contrast, the top performing challenge participant, submitted by a team who took part in a 2-week coding and genetics bootcamp, was a minimalist random forest meta-meta predictor trained on a personal laptop. It should also be noted that AlphaMissense was published after the %WT activity values were publicly released, though these data were not directly used to train their model. Despite the disparity in resources needed to train each model, AlphaMissense only performed marginally better than Bootcamp Team 3’s model.

Determining the impact of missense variation on protein function and pathogenicity remains a complex challenge. Given the influence of VUS on the diagnostic odyssey, advances in the field have the potential to directly impact patients (Bauskis et al, 2022). As more VUS are encountered in newborn screening due to increasing availability of treatments for rare diseases, predictors of variant functional effect will be valuable tools for quickly interpreting their potential relevance to disease (Stark and Scott, 2023). This is especially true for MLD, which has just seen the approval of new therapeutic approaches, and for which early diagnosis is essential for effective intervention (Adang et al, 2024). While the field of variant effect prediction has yet to see advances in performance seen in the field of protein structure prediction (Kryshtafovych et al, 2021), top models incorporate information from state-of-the-art structure prediction tools (Cheng et al, 2023). In many aspects protein structure prediction may be an easier learning task than predicting a mutation’s pathogenicity. It is possible to determine the position of a molecule in a crystallized protein structure at angstrom resolution, whereas the gold standard for determining the disease relevance of a rare missense variant requires observing it multiple times in patients (Richards et al, 2015).

While MLD is a monogenic disease, whose severity can be largely explained by the residual activity of two enzymes, ARSA and PSAP, many human diseases are polygenic, or have strong polygenic backgrounds that can influence the penetrance of mutations in genes associated with monogenic disease (Khan et al, 2023). Studies of genes such as *ARSA*, which have clear functional readouts, can help train better computational models. This will consequently have the potential to improve predictions for genes that are harder to study, advancing the field of clinical and medical genetics.

## Supplementary information

**Fig. S1**: **Model performance based on key metrics:** Model performance based on Pearson’s correlation, Kendall’s tau, AUC and Truncated AUC. The best-performing model for each team is shown in blue. Baseline models and individual feature (evolutionary and structure) based performance are shown in grey. Models are sorted on the y-axis based on their average ranking according to the four metrics used.

**Table S1**: **Mutation Information:** A list of ARSA missense mutations from (Trinidad et al, 2023), their observed %WT activity values, and submitted scores from CAGI challenge participants, supplementary publicly available models, and fundamental features. An indicator column (*evaluationSet*) denotes whether variants were included in the final evaluation dataset.

**Table S2**: **Model performance based on key metrics:** Performance of top methods from each team according to the evaluation metrics described in Section 3.2, and their respective rank according to each metric.

**Table S3**: **Difficult to predict pathogenic mutations**. A table of difficult-to-predict pathogenic mutations, generated as described in Section 3.6.

**Table S4**: **Difficult to predict benign mutations**. A table of difficult-to-predict benign mutations, generated as described in Section 3.6.

**Table S5**: **ID mapping:** A mapping between raw model ID’s and those used for each team’s top performing model in figures.

## Acknowledgments

Authors would like to acknowledge Evan Witt, Nick Knobloch, Raymund Bueno, and Larry Hengl for volunteering as instructors in the ARSA CAGI Bootcamp.

## Declarations

- Funding The CAGI experiment was supported in part by the National Institutes of Health (NIH) awards U24HG007346 (SEB), U01HG012022 (PR), R35GM124952 (YS), R01HG012117 (EF), AG068214 (OL), and Spanish Minesterio de Ciencia e Innocavion grants PID2019-11217RB-I00 and PID2022-142753OB-I00 (XC).
- Conflict of interest/Competing interests Wyatt T. Clark, Marena Trinidad, Courtney Astore, Teague Sterling, and Sufyan Kazi are former employees and potential shareholders of BioMarin Pharmaceutical. Rocio Acuna-Hidalgo is a current employee and shareholder of Nostos Genomics GmbH. Andrea Grafmüller and Laura T. Jiménez Barrón are former employees of Nostos Genomics GmbH.
- Ethics approval Not relevant.
- Consent to participate Not relevant.
- Consent for publication All authors have consented to publication of this manuscript.
- Availability of data and materials All data are available as supplementary files, or as supplementary files from Trinidad et al (2023).
- Code availability Not relevant.
- Authors’ contributions SJ, MT, KL, SL, PR, TB, WTC contributed to manuscript preparation. MT, SJ, CA, AK, VP, RR, MV, DZ, RM, TS, JG, JLM, SK, SEB, TB, PR, DS, TS, and WTC were bootcamp organizers or instructors. TBN, CB, SDN, ED, KJ, FG, AG, CJ, GM, BW, KR, SZC, LC, SB, BO, and FO were bootcamp participants. TBN, CB, SDN, ED, KJ, FG, AG, CJ, GM, BW, KR, SZC, LC, SB, BO, FO, KC, PK, AW, OL, SR, SP, RS, RS, DJ, EF, RJ, AK, JX, ZS, SO, NP, XC, RAH, AG, LJB, MM, CS, GB, PLM, RC, YS, SZ, YS, FP, MR, GC, DR, PH, SK, and EC were ARSA CAGI challenge participants. MT and WTC were ARSA CAGI challenge data providers. SJ, KL, SL, TB, PR, WTC, and SEB were ARSA CAGI challenge assessors or organizers.

## Notes

### Summary of Updates

Fixed a typo in an author's name.

