## Supplementary Figure 1 for "Evaluation of enzyme activity predictions for variants of unknown significance in Arylsulfatase A"

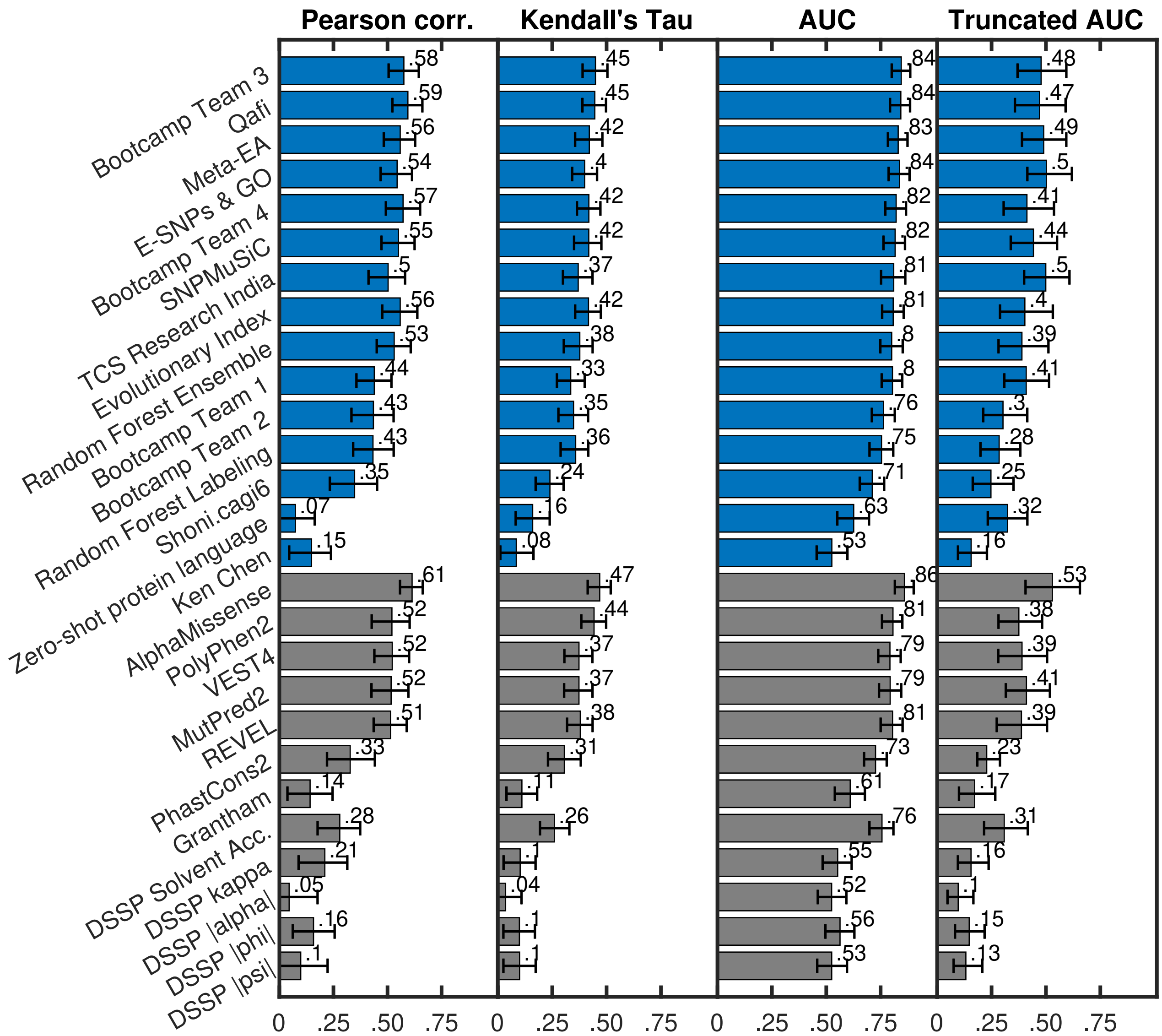


Figure S1: Model performance based on key metrics: Model performance based on Pearson's correlation, Kendall's tau, AUC and Truncated AUC. The best-performing model for each team is shown in blue. Baseline models and individual feature (evolutionary and structure) based performance are shown in grey. Within each group (submitted models, baselines, evolutionary and structural features), models are sorted on the y-axis based on their average ranking according to the four metrics used. For teams that submitted multiple models, we show the performance of their best model.
